# RegEnrich: An R package for gene regulator enrichment analysis reveals key role of ETS transcription factor family in interferon signaling

**DOI:** 10.1101/2021.01.24.428029

**Authors:** Weiyang Tao, Timothy R.D.J. Radstake, Aridaman Pandit

## Abstract

Changes in a few key transcriptional regulators can lead to different biological states, including cell activation and differentiation, and diseases. Extracting the key gene regulators governing a biological state allows us to gain mechanistic insights and can further help in translational research. Most current tools perform pathway/GO enrichment analysis to identify key genes and regulators but tend to overlook the regulatory interactions between genes and proteins. Here we present *RegEnrich*, an open-source Bioconductor R package, which combines differential expression analysis, data-driven gene regulatory network inference, enrichment analysis, and gene regulator ranking to identify key regulators using gene/protein expression profiling data. By benchmarking using multiple gene expression datasets of gene silencing studies, we found that *RegEnrich* using the GSEA method to rank the regulators performed the best to retrieve the key regulators. Further, *RegEnrich* was applied to 21 publicly available datasets on *in vitro* interferon-stimulation of different cell types. We found that not only IRF and STAT transcription factor families played an important role in cells responding to IFN, but also several ETS transcription factor family members, such as ELF1 and ETV7, are highly associated with IFN stimulations. Collectively, *RegEnrich* can accurately identify key gene regulators from the cells under different biological states in a data-driven manner, which can be valuable in mechanistically studying cell differentiation, cell response to drug stimulation, disease development, and ultimately drug development.

## 1. Introduction

The advances in high throughput technologies such as genomics, transcriptomics, and proteomics have provided unprecedented opportunities to mechanistically understand the genetic and epigenetic alterations in diseases, cellular development, cell stimulation, and immune activation ^1,2^. Typically, alterations in the expression of key genes and proteins play a central role in many of these biological states. Thus, to understand the differences between these states, one of the fundamental steps is to identify a set of genes or proteins that are differentially expressed ^3^. To understand the underlying biological process and functions of these molecules, annotation enrichment methods, such as pathway and gene ontology (GO) term enrichment, have been used widely ^4^. Although the enrichment analysis provides crucial clues about the underlying biological processes and pathways, lack of information about the underlying regulation hinders us to mechanistically understand how these biological states can be achieved.

To study the function, regulation and dynamics of individual genes (or proteins) in a complex biological system, network biology is emerging as an important tool ^5^. Several studies have demonstrated that constructing gene/protein interaction networks allows us to gain important insights into the regulatory mechanisms that govern different biological states including disease, cellular activation, and differentiation ^6,7^. In a gene/protein interaction network, densely connected genes (or *hub genes*) have been shown to be crucial for network’s integrity and the corresponding biological state ^6,7^. However, considering only topological parameters (such as hubness or degree) of a network may overlook key regulators ^5^. So, to gain regulatory insights, we should consider both network topology and the corresponding alterations in gene or protein expression.

Transcription factors and co-factors (TFs) can directly (and/or indirectly) regulate the expression of multiple target (and/or downstream) genes and proteins ^8–10^. Some studies took advantage of curated TF–target networks, and used Fisher’s exact test to analyze the functional enrichment of the target genes ^11^. However, current curated networks are incomplete, and increasing studies have shown that regulatory interactions may differ over time, upon different conditions and cellular states in the same organism ^8,12,13^. So, analyses based on these incomplete static networks might not be sufficient to unveil functional regulatory patterns in complex biological processes.

State/cell/condition specific gene regulatory network can directly be inferred from the gene or protein expression data (data-driven network) ^8,14^. Using these data-driven networks and results from differential expression analyses, one can deduce key regulators. For example, ARACNE and ARACNe-AP, which have been used in reverse engineering field, reconstruct gene regulatory network from gene expression profile datasets based on mutual information ^15,16^. The VIPER R package takes advantage of this network and use t-statistics by comparing gene expression of different conditions to compute the final enrichment p-values for TFs ^17^. However, since the VIPER package calculates the final enrichment p-values using t-statistics, it is currently not applicable for users to apply this tool to identify the key regulators in other experimental settings, such as time series experiments.

To address aforementioned problems, we developed “*RegEnrich*”, an open-source R package for gene regulator enrichment analysis (**Fig. 1**). The aim of *RegEnrich* pipeline is to identify the key regulators based on their differential expression and enrichment of their potential downstream targets from a given gene set. Because the gene regulators do not act alone but function as part of a complex network, by using *RegEnrich* one can refine a key gene regulatory network to study the biological process and visualize the derived network.

**Fig. 1.**
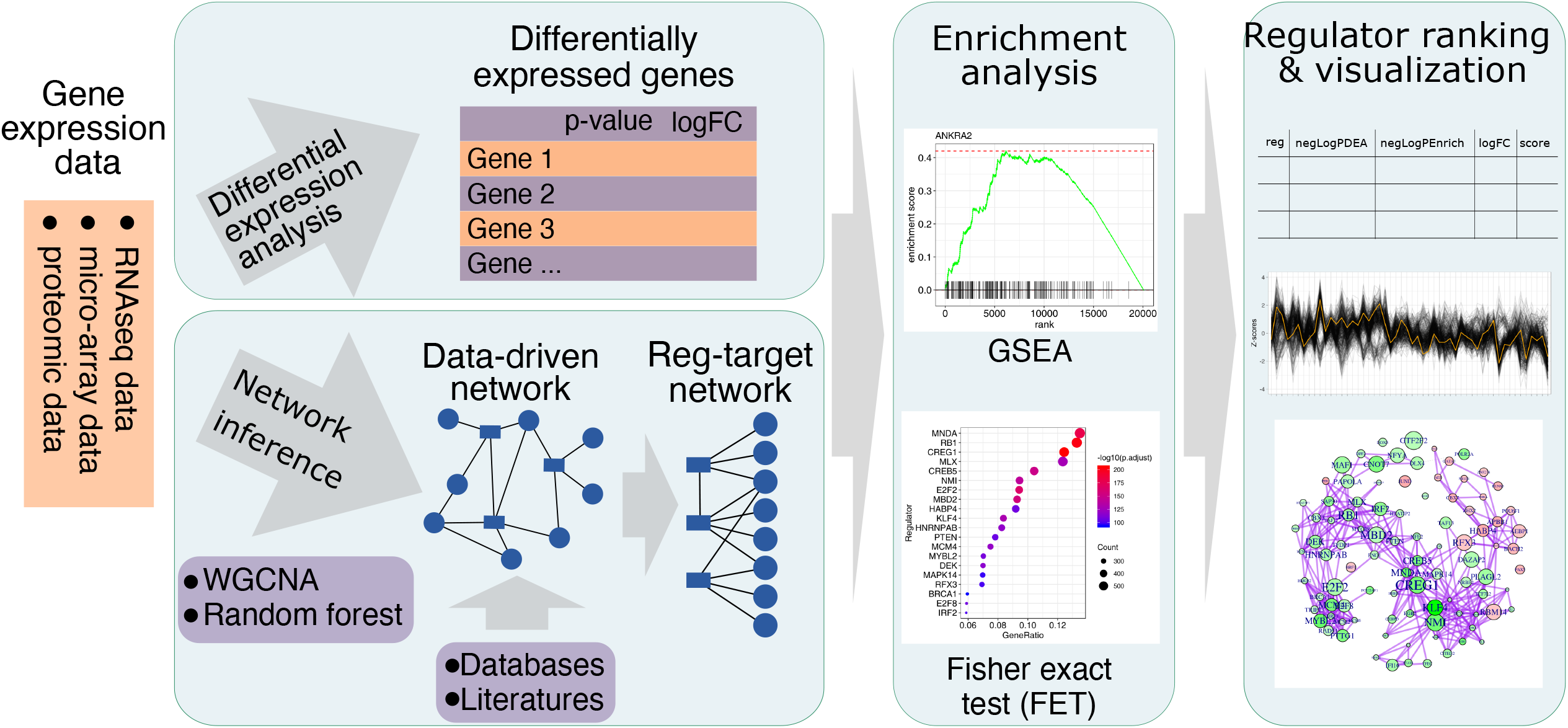
The analytic workflow of *RegEnrich* package. *RegEnrich* consists of four major steps: differential expression analysis, regulator-target network construction, enrichment analysis, and regulator ranking and visualization.

## 2. Methods

The *RegEnrich* is a modular pipeline and consists of four major steps: (a) differential expression analysis; (b) regulator-target network construction; (c) enrichment analysis; and (d) regulator ranking and visualization.

### 2.1 Differential expression analysis

*RegEnrich* pipeline can be applied to multiple gene expression datasets, including RNA sequencing (RNAseq), micro-array and proteomic data. The first step of finding the key regulators is to obtain differentially expressed genes or proteins (DEs), and corresponding differential significance p-values (P_D_) and fold changes between conditions. With respect to two-group comparison, here, *RegEnrich* incorporates Wald significance test from DESeq2 package and the empirical Bayes method-based linear modelling from limma package to perform the differential expression analysis on RNAseq data and micro-array/proteomic data, respectively ^18,19^. Regarding multiple groups comparisons and more complex experiment settings such as time series study, the negative binomial generalized linear model (GLM)-based likelihood ratio test (LRT) from DESeq2 package ^18^ and linear model-based LRT are implemented for RNAseq data and micro-array/proteomic data, respectively.

### 2.2 Regulator-target networks inference

There are two major types of gene regulatory networks (or regulator-target networks) being proposed: static network and dynamic network ^20–22^. In a static network, genes are expressed in a steady state thus cannot describe the dynamics of an evolving process, while genes are dynamical in a dynamic network ^23^. These networks can be constructed by many different computational approaches ^14,15,24–27^. Here, the regulator-target network inference is based on four hypothesis: (1) the gene regulatory network is a snapshot of a dynamic network within the users’ experiments; (2) It is a directed network, where the edges are from a regulator to its targets, or from a regulator to its targeted regulators; (3) the potential regulators are transcription factors and co-factors (this can be changed in *RegEnrich* by users); and (4) the expression change of a regulator can lead to the expression change of its downstream targets. Here, the targets are not only direct targets that the regulator binds to but also the downstream genes whose expression can be perturbed by the regulator. Presently, *RegEnrich* provides users two basic options to infer regulator-target network, i.e., COEN (co-expression network) and GRN which is based on random forest algorithm.

#### COEN

Here, the co-expression network is constructed based on WGCNA (weighted gene co-expression network analysis) algorithm ^25^. And it can be summarized as three major procedures. Firstly, similarity matrix is calculated using correlations in expression data to measure the relationship strength between each pair of genes (nodes). Secondly, by applying the approximate scale-free topology criterion, raising the co-expression similarity to a power to define the weighted network adjacency matrix. Thirdly, this adjacency matrix is then used to calculate the topological overlap measure (TOM), which reflects not only the similarity of each pair of nodes but also their neighbors’ similarity ^6^. The TOM defines the final co-expression network ^25^.

#### GRN

This ensemble regression tree-based method was initially described in GENIE3, which was the best performer in the DREAM4 *In Silico Multifactorial* challenge ^14^. The basic idea of GENIE3 is that each gene is regressed in turn against all other genes in order to obtain network weights (edge weights), which quantify the strength of the dependence of each pair of genes. The edge weight (*W_ij_*) is the importance of gene i in the tree model predicting gene j, which can be interpreted as the fraction of variance of the expression of gene j that can be explained by gene i ^27^. However, the GENIE3 package is slow especially when it is deployed on genome-wide studies with a large number of experiments. So, to facilitate usage and improve speed, we implemented this algorithm by allowing users to define their own regulators and by supporting parallel computing (**Fig. 2**). In addition, in this random forest-based method we found the expressions of some genes are hardly predicted by other genes. So, we modified this algorithm by adding a filtering procedure to remove the poor random forest models (**Fig. 2**). In other words, this procedure removes the genes, and corresponding edges from the final network, whose expression is hardly predicted by the expression of the predefined regulators.

**Fig. 2.**
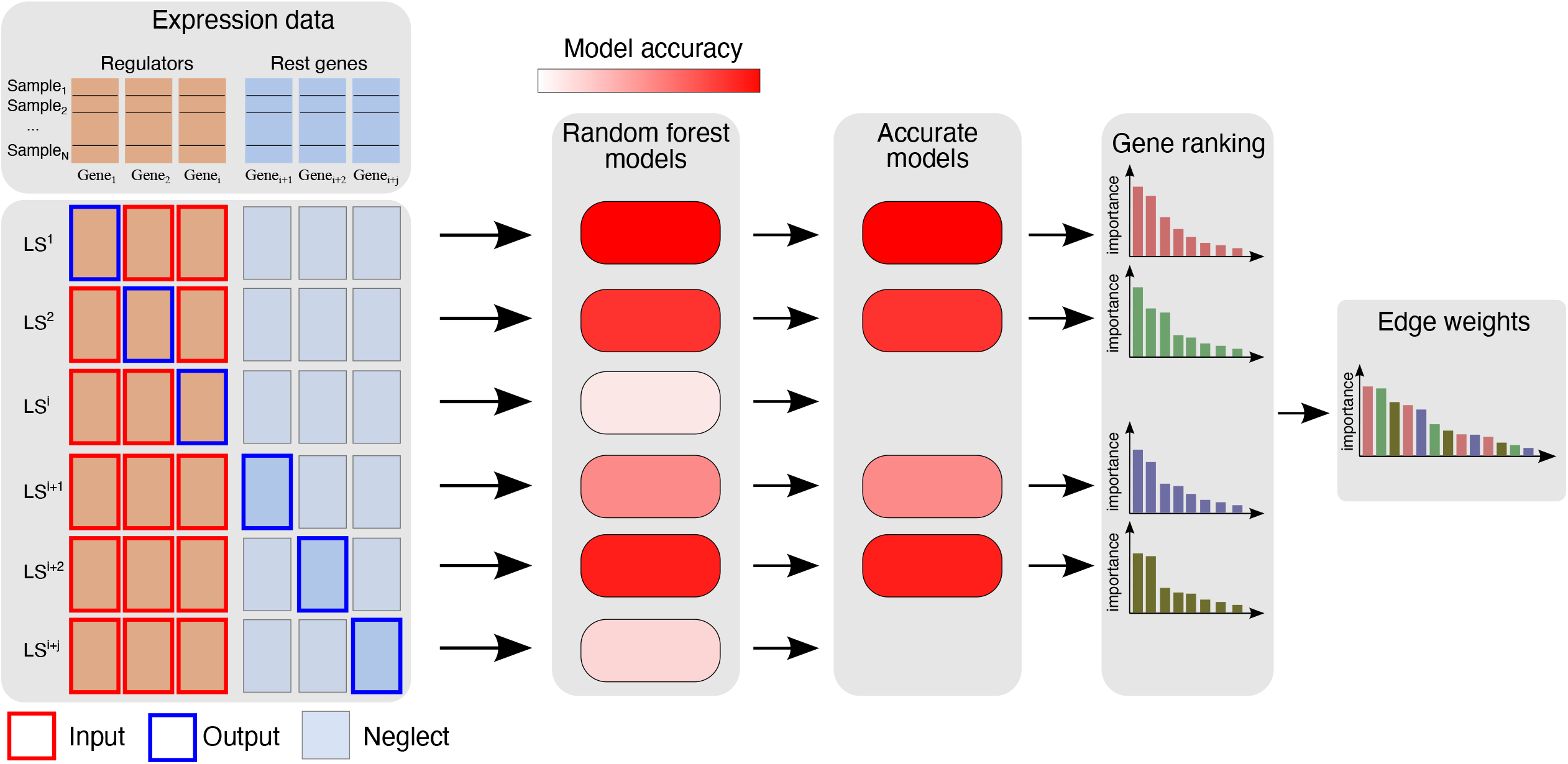
Fig.2. The workflow to construct a random forest-based network. According to the regulators predefined by the users, the gene/protein expression matrix split up into two parts, i.e., the expression of regulator genes and the expression of non-regulator genes. For each regulator gene r = 1,…,i, a learning sample LS_r_ is generated with expression levels of r as output values and expression levels of all other regulator genes as input values. A random forest model m_r_ is fitted from LS_r_ and local weights of all regulator genes except r are computed. While for each non-regulator gene n = i+1,…,i+j, a learning sample LS_n_ is generated with expression levels of n as output values and expression levels of all regulator genes as input values. A model m_n_ is fitted from LS_n_ and local weights of all regulator genes are computed. The model m_r_ or m_n_ is removed if the model accuracy, according to out-of-bag accuracy, is too low. Finally, all of the local weights in accurate models are then aggregated to get the global weights of network edges. Note that when users construct the network by setting all genes as the regulators, the minimum accuracy as the threshold of model accuracy and using one computer core, this method is equivalent to GENIE3.

We provide users an option to either supply a list of regulators of their interest or use the default list of regulators provided in *RegEnrich*, which were retrieved from three studies ^28–30^. Using either COEN or GRN network, we then extract the regulator-target network, by retaining the top ranked edges (default = top 5% edges) between the regulators and their targets, and subsequently filtering out non-connected nodes. Apart from the data-driven network, *RegEnrich* also allows users to provide their own regulator-target network, which can be derived from the literature, databases or defined by the user using their own data of other types.

### 2.3 Enrichment analysis

The regulators are considered key regulators if they are differentially expressed along with their own targets in a differentially expressed gene set. In other words, not only these regulator genes but also their target genes are differentially expressed upon different conditions. Finding these key regulators is an enrichment task, which is similar to retrieving the most overrepresented (enriched) biological annotations, such as gene ontology and pathways terms, of a list of interesting genes. Presently, *RegEnrich* provides users two options: Fisher’s exact test (FET) and gene set enrichment analysis (GSEA).

<u>Fisher’s exact test (FET)</u>, also known as hypergeometric test, calculates probability using the hypergeometric distribution (Equation 1). This distribution describes the probability of the number of draws being successful (*k*) within a sequence of draws (*M*), without replacement, from a finite population (*N*) consisting of two types of elements (the total number of successful types is *s*).

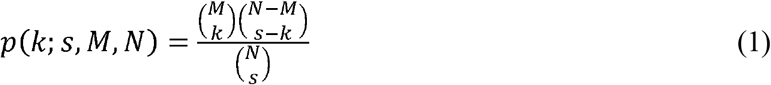

Then the p value, depicting the probability of observing K (and more) differential targets by chance, of regulator *i* being overrepresented, is calculated by

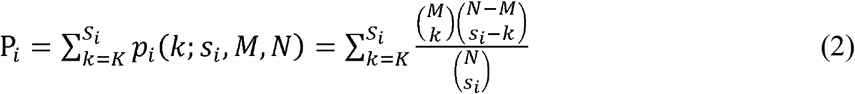

where N is the total number of genes in the previously constructed regulator-target network; M is the number of genes in the list of users’ interests (the genes not in the network are excluded), which is typically the differential genes between conditions; *s_i_* is the number of target genes of regulator *i* in the network; K is the number of target genes that are also in the list of users’ interests. This process repeats for all regulators that are predefined by users.

Gene set enrichment analysis <u>(GSEA)</u> is one of the most widely used methods to study biological function of groups of genes and to interpret gene expression data ^31^. GSEA takes into account all of the genes in an experiment, unlike FET that takes into account only those genes above a fold-change or significance cutoff. Here, *RegEnrich* takes two basic inputs, the TF-target network and a named vector of decreasingly sorted ranking metrics (r, z-score scaled negative logarithm of differential significance p values) of all genes. Briefly, there are three major steps in this analysis:

1. Calculation of an Enrichment Score (ES) by:

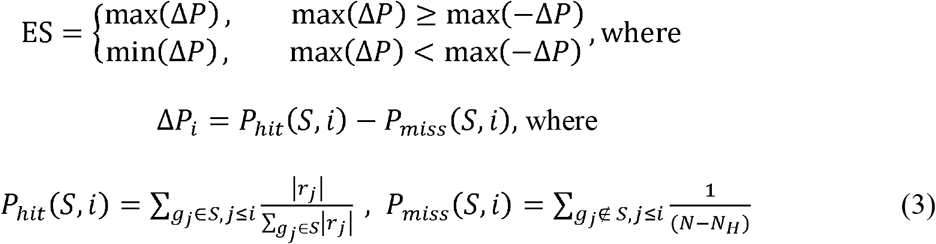

Here, *i* is the index of decreasingly sorted ranking metrics *r*, *S* is the set of target genes of one particular regulator, *N_H_* is the number of genes in *S*, and *N* is the total number of valid genes in the regulator-target network.
2. Randomly shuffle the ranking metrics of genes, and re-compute ES. And repeat this process for 1,000 permutations to generate ES_NULL_ that establishes an empirical distribution. Estimate empirical p value for S from ES_NULL_ by only positive portion of the distribution corresponding to the sign of the observed ES.
3. Perform (1) and (2) for each regulator, resulting a numeric vector (*P_E_*) in which each value is an enrichment p value for each regulator.

### 2.4 Regulator ranking and visualization

After the enrichment analysis by either FET or GSEA, the overall ranking scores of regulators were calculated by:

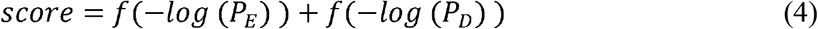

where 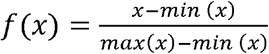, and *P_D_* is the vector of p-values of regulators obtained from differential expression analysis, *P_E_* is the vector of p-values of regulators obtained from the enrichment analysis.

In the *RegEnrich* package, we have implemented several functions for visualizing the information of regulator and its targets (**Fig. 1**). For example, “plotRegTarExpr” function is to plot the expression pattern of regulator and its targets.

## 3. Results

### 3.1 Time consumption and memory usage by *RegEnrich*

The most time- and memory-consuming procedure of the *RegEnrich* pipeline is inferencing a regulator-target network from the gene expression data. Here, we benchmarked the time consumption and memory usage of different methods in the *RegEnrich* package using Intel^®^ Xeon^®^ Processors with one (**Fig. 3**) or four cores (**Fig. S1**) on CentOS Linux 7 operating system on high-performance computing facility at University Medical Center Utrecht. And the gene expression data were simulated with different numbers of samples (10, 20, 50, 100 for COEN method and 50, 100, 200 for random forest method (GRN)) and different number of genes (from 2,000 to 40,000). Overall, the speed of both methods decreased with the increase of the number of genes, and the speed was also dependent on the sample size for only the GRN method (**Fig. 3A**). More specifically, the consumed time of the COEN method increased quadratically with the number of genes, while independent on the sample size. The COEN method was around 1 ~ 100 times faster, compared to random forest method, when the number of genes was below 20,000 and the number of samples was over 50. However, since the random forest method is linearly, rather than quadratically, dependent on the number of genes, The COEN method spent more time when the number of genes was above 25,000 and the sample size was below 100. Network construction using the GRN method running on 4 CPU cores was about 2 times faster than the single-threading implementation under all circumstances (**Table S1**).

**Fig. 3.**
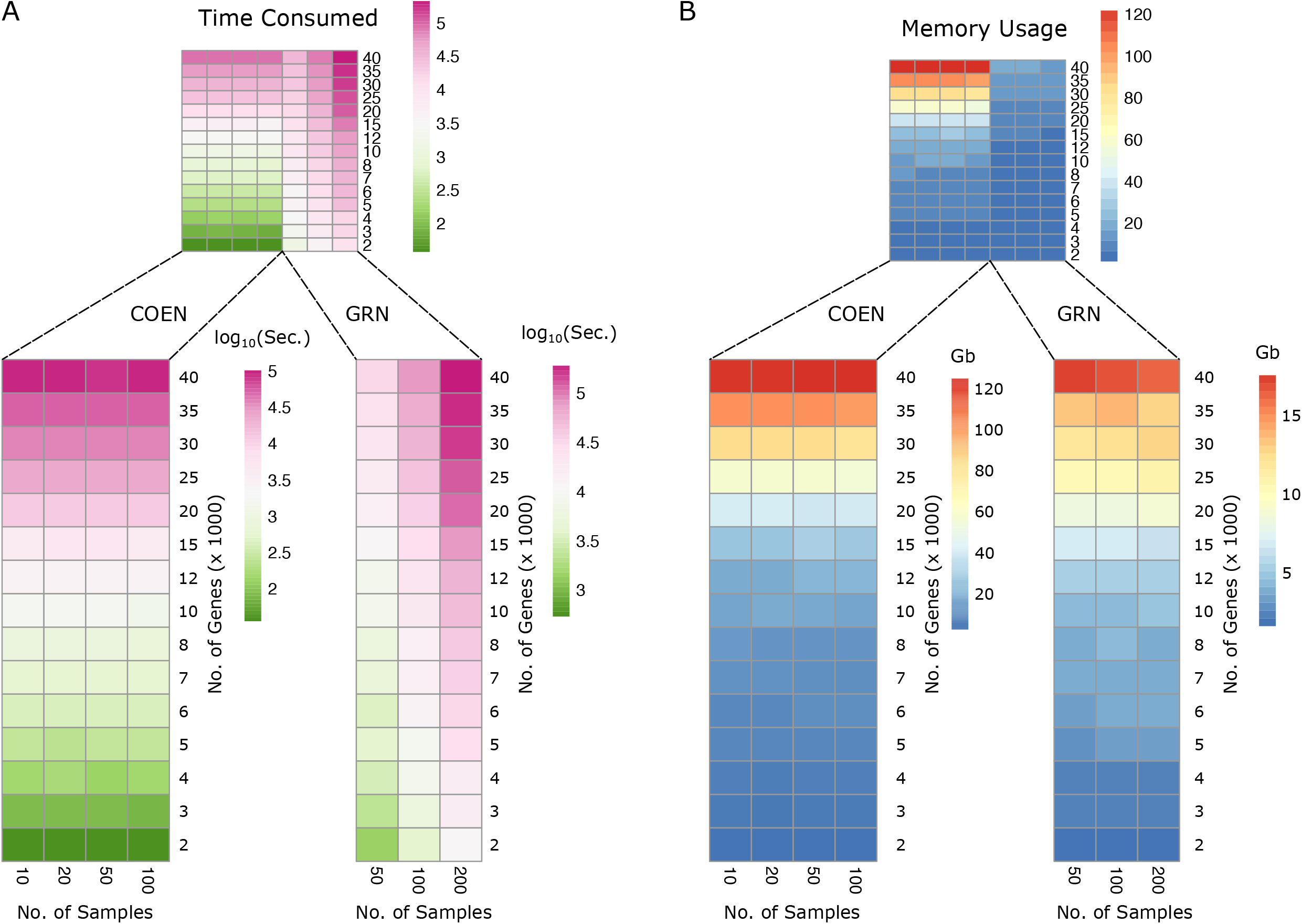
Time consumption and memory usage by *RegEnrich*. With one CPU core, (**A**) the time consumed, and (**B**) maximum memory used, by *RegEnrich* when analyzing a gene expression dataset with different number of genes (ranging from 2,000 to 40,000) and different number of samples (ranging from 10 to 100 and from 50 to 200 for COEN and GRN network, respectively).

The maximum memory usage of the COEN method again increased quadratically with the increase of number of genes, and independent with the number of samples (**Fig. 3B**). Given the same size of the simulated expression data, the memory used by the COEN method were more than that by the GRN method in almost all circumstances. In general, using a dataset of 50 samples based on 1 CPU core, when constructing a protein coding network, which comprises about 2×10^3^ genes, it costs ~ 3.4 hours (38.5 Gb) and ~ 4.1 hours (8.2 Gb) by the COEN and by the GRN method, respectively. And it costs 1.6 hours (7.8 Gb) by the GRN method using 4 CPU cores. The figure may be different depending on the computing power of the processors, but the order of magnitude will less likely change. So, the users can roughly expect the time and memory usage when performing the *RegEnrich* analysis according to **Fig. 3** and **Table S1**.

### 3.2 Comparisons of key regulators obtained by different methods

Increasing studies predict key gene regulators by the hubs in a network, which is defined by topological features, such as degree and closeness centrality ^32,33^. The degree of a node is the total number of nodes connected to this node in a network. The out-degree of a node is the number of nodes pointed by this node in a directed network. The closeness of a node is defined as the reciprocal of the sum of the shortest path length between this node and all other nodes in the network ^34^. And the out-closeness of a node is defined as the reciprocal of the sum of the shortest path length from this node to all other nodes in a directed network. We applied out-degree, out-closeness, and *RegEnrich* on a paired transcriptomic dataset of patients with Lyme disease (paired gene expression samples from 26 donors on two time points) ^35^ to identify hubs (or key regulators) in the network. As a result, we obtained three sets of hubs defined by 50 regulators with highest out-degree (degree hubs), out-closeness (closeness hubs) or *RegEnrich* ranking score (*RegEnrich* key regulators). We found 20% (10) regulators were overlapping between degree hubs and closeness hubs, while *RegEnrich* key regulators were barely overlapping with degree hubs or closeness hubs (**Fig. 4A**). Similarly, more overlapping regulators were observed between degree hubs and closeness hubs in the network constructed using ARACNE-AP package, and these hubs are rarely the key regulators identified by VIPER package (**Fig. 4B**). Different network-inferencing methods and enrichment methods within *RegEnrich* showed consistent top ranked regulators (**Fig. 4C and D**), which are overlapping nearly half of the regulators identified by VIPER (**Fig. 4C**). Altogether, both *RegEnrich* and VIPER tend to rank the non-hub regulators to be the key regulators. These results might suggest that the hub regulators with high out-degree or high out-closeness are too important in the survival-related biological processes to be strongly perturbed, thus may not play central roles in the biological process being currently studied.

**Fig. 4.**
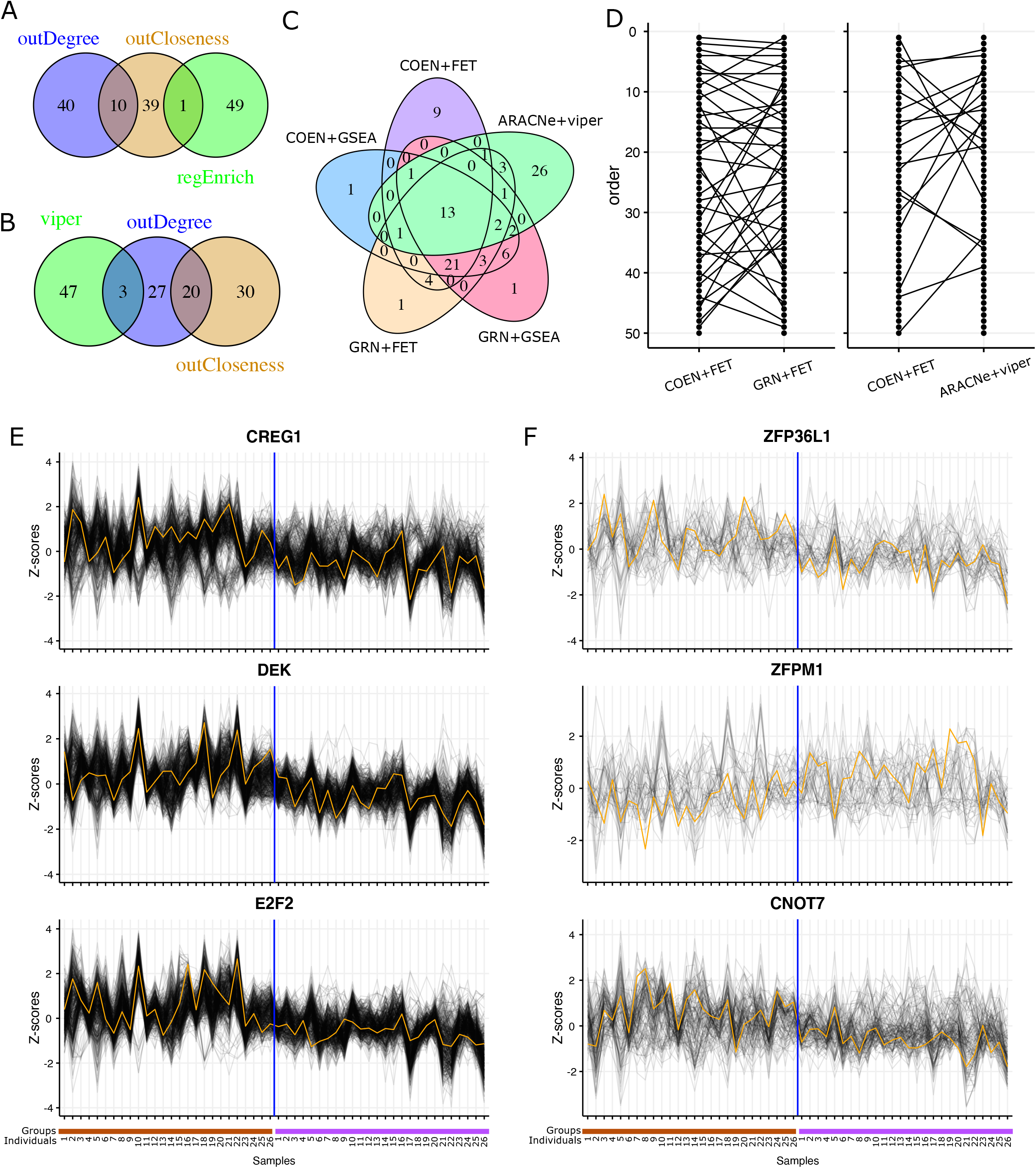
Comparison of key regulators (or hubs) identified by different methods using the data obtained from [^35^]. Venn diagram shows the overlap between the top 50 hubs/regulators **(A)** defined by out degree, out closeness and the *RegEnrich* score (using COEN network and FET enrichment method); **(B)** defined by out degree, out closeness and the VIPER package (using ARACNe network). **(C)** Venn diagram shows the overlap between the top 50 hubs/regulators defined by *RegEnrich* (using different parameter combinations) and those by VIPER package. **(D)** Ladder plots compare the rank of regulators identified by (left) *RegEnrich* using different network inferencing methods, and those (right) by *RegEnrich* and by VIPER. Lines connect the same regulators. The expression pattern of top three key regulators and their targets identified **(E)** by *RegEnrich* using COEN network and FET method and by **(F)** VIPER. Orange lines are the normalized expression of regulators and grey lines are that of the targets of the regulators. The brown bars on the x-axis indicate samples at week 1, and purple bars the samples at week 3.

Although a decent number of regulators were commonly ranked in the top 50 regulators by both *RegEnrich* and VIPER, most of top10 regulators identified by these two packages are unique (**Fig 4D** right). Clearly, the expression of the 3 topmost-ranked regulators (CREG1, DEK, and E2F2) identified by *RegEnrich* were differential between week 1 and week 3, and shows very similar pattern to that of corresponding targets (**Fig. 4E**). Although the expression of 3 top-ranked regulators identified by VIPER was also differential, only the expression of the targets of CNOT7 were correlated with that of CNOT7 itself (**Fig. 4F**). One may argue that it is inevitable that the expression of a regulator and its targets are highly correlated in COEN networks. This correlation may not hold true for the network that was built by ARACNE-AP using mutual information. However, such correlations between regulator and its targets were still observed when the network is build using random forest, which is also non-linear method (**Fig. S2**). This implies that *RegEnrich* might identify the key regulators that both their own expression and their targets’ expression associates with the biological process of interest.

### 3.3 *RegEnrich* is able to identify the key regulators in gene silencing studies

RNA interference (RNAi)-mediated gene silencing has been widely used to study the biological function of the silenced gene. In a gene regulator silencing experiment, the successfully silenced gene is typically the key regulator that has been malfunctioned. The performance metrics of *RegEnrich* and VIPER were compared by the ability to rank the silenced gene as one of the top key regulators. Here nine gene silencing experiments from four independent datasets ^36–39^ were used to benchmark *RegEnrich* using COEN network with either FET or GSEA enrichment methods, and VIPER using the network built by the ARACNE-AP package. Multiple cell lines/types, different silenced genes, and varied number of samples were deliberately included in these datasets to evaluate the bias induced by these variables (**Table 1**). *RegEnrich* with either FET or GSEA enrichment method outperformed VIPER in all these datasets; and within *RegEnrich*, GSEA outperformed FET in most cases. For example, using GSEA method, STAT3 and FOXM1 were ranked as the top key regulators when STAT3 and FOXM1 were silenced in the BTIC and ST486 cell line, respectively (**Table 1**). STAT3 and FOXM1 were also ranked high (the second key regulators) in these experiments when we applied *RegEnrich* with the FET method was applied. Interestingly, in GSE17172 dataset, although FOXM1 was not ranked as the first regulator using *RegEnrich* FET, FOXN3 and FOXG1, other two genes from the same FOX transcription factor family were ranked as the first and fourth regulator, respectively (**Table S2**). This implies that at least several members of the FOX family were perturbed by FOXM1 silencing due to either off-targeting or downstream transcriptional signaling and can be inferred by *RegEnrich*. By applying VIPER to this dataset, STAT3 was ranked as the 7^th^ regulators, however, FOXM1 was failed to be identified by using VIPER maybe because the sample size is small (**Table 1**).

**Table 1.** The transcription regulators identified by *RegEnrich* in gene silencing studies*

| GEO accession | Silencing technology | No. of samples | Cell line | Silenced gene(s) | Ranking |  | ARACNE + VIPER |
| --- | --- | --- | --- | --- | --- | --- | --- |
|  |  |  |  |  | *RegEnrich* (FET) | *RegEnrich* (GSEA) |  |
| GSE19114 | shRNA | 44 | BTIC | STAT3 | **2** | 1 | 7 |
| GSE17172 | shRNA | 9 | ST486 | FOXM1 | **2** | 1 | - |
| GSE19114 | shRNA | 12 | SNB19 | STAT3 | **14** | 5 | n/a |
| GSE2350 | siRNA | 8 | Burkitt lymphoma cell line | BCL6 | **32** | 11 | - |
| GSE19114 | shRNA | 12 | SNB19 | CEBPB | **50** | 24 | n/a |
| GSE19114 | shRNA | 44 | BTIC | STAT3 & CEBPB | **2** & 285 | 1 & 957 | 6 & n/a |
| GSE19114 | shRNA | 12 | SNB19 | STAT3 & CEBPB | 365 & **6** | 38 & 11 | n/a & n/a |
| GSE51978 | shRNA | 9 | IMR32 | CHAF1A | **10** & 29<sup>#</sup> | 15 & 46<sup>#</sup> | - |
| GSE19114 | shRNA | 44 | BTIC | CEBPB | 913 | 793 | n/a |

\* “n/a” means no result for the regulators of interest available after ranking procedure. “-” indicates that
ARACNE failed to construct a network based on the dataset. <sup>#</sup> means the ranking on day 5 and day 10 according to the experimental setting, respectively.

Similarly, STAT3 and CEBPB in SNB19 cell line, CHAF1A in IMR32 cell line, and BCL6 in Burkitt lymphoma cell line were identified by *RegEnrich*, with GSEA method, as one of top 20 key regulators in each corresponding dataset. Of note, the rankings of these regulators were all still high because these are regulators popping up from a total list of 1712 regulators in these *RegEnrich* analyses. Meanwhile, we also assessed the *RegEnrich* using two datasets, where STAT3 and CEBPB were tried to be simultaneously silenced in either BTIC or SNB19 cell lines. Even though these two genes were intended to be silenced, only one gene was successfully silenced, for example STAT3 gene in BTIC cell line and CEBPB gene in SNB19 cell line. As expected, all approaches top-ranked only STAT3 gene in BTIC cell line, while only *RegEnrich* able to top-ranked CEBPB gene in BTIC cell line (**Table 1**). To evaluate *RegEnrich*’s ability of filtering the false positive results, we included a dataset, where CEBPB was not successfully silenced in BTIC cell line. All three approaches did not rank CEBPB as one of the top regulators. Altogether, *RegEnrich* with COEN network and GSEA method is robust to identify the key regulators in well controlled *in vitro* experiments even when the sample size is small.

### 3.4 *RegEnrich* retrieves interferon related regulators

In human there are three types of interferons (IFN): type I IFNs (IFNα, β, ε, κ, and ω); type II IFN (only IFNγ); and type III IFNs (IFNλ1, λ2, λ3, and λ4) ^40,41^. Due to a great therapeutic value of IFNs against virus infection and cancer, multiple studies have been performed to study the regulatory mechanisms of IFNs and interferon-stimulated genes (ISGs). For example, it has been revealed that extracellular IFNs activate cells by a signal transduction cascade, including activating transcription factors STATs and/or IRFs leading to the induction of hundreds of ISGs, and forming a frontline of defense against virus infections ^40,41^. However, the mechanisms underlying the regulation of most of these ISGs may vary between different cell types and tissues, and remains incompletely understood.

Given the potential of identification of key regulators by *RegEnrich* in a biological process, we tried to predict the key regulators, by which IFNs stimulated cells to express ISGs. We retrieved and analyzed 11 microarray or RNAseq datasets from GEO database, comprising 21 *in vitro* experiments, in which different cells were stimulated by either type I or type II IFN (**Table 2**). We found that *RegEnrich* identified STAT transcription factor family members, including STAT1 and STAT2, in most IFN stimulation experiments, which is supported by the well-known IFN signaling pathway ^41^. In addition, IRF (interferon regulatory factors) transcription factor family members, such as IRF9 and/or IRF7, were also identified as key regulators in majority of the type I IFN stimulation experiments (**Table 2**). These IRFs have been reported to play important roles in producing of type I IFN downstream receptors that detect viral RNA and DNA, and in regulating of interferon-driven gene expression ^42^.

**Table 2.**
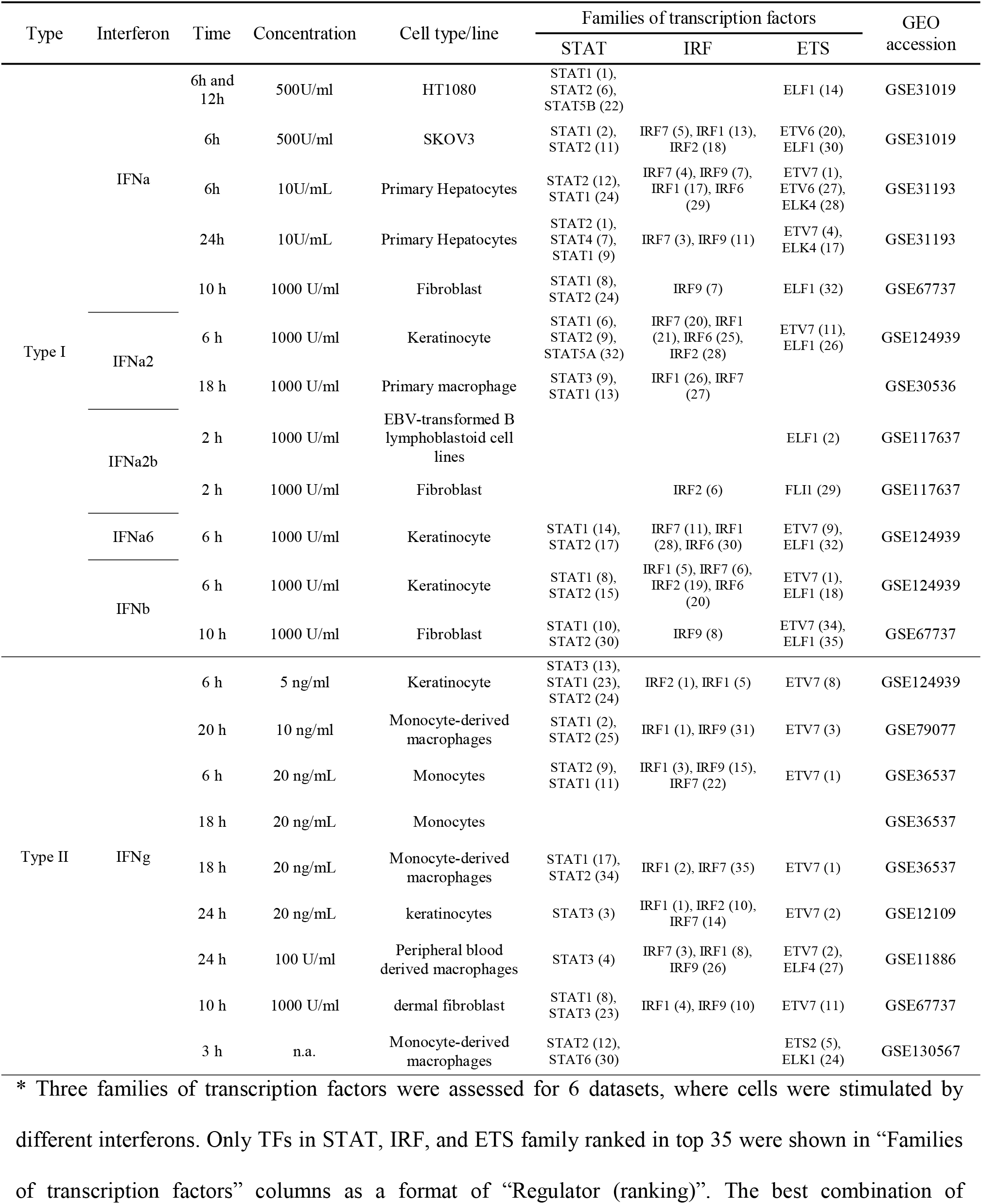
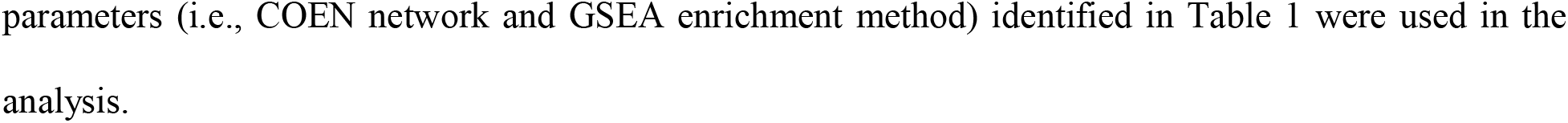
The transcription factors identified by *RegEnrich* in interferon stimulation studies*

| Type | Interferon | Time | Concentration | Cell type/line | Families of transcription factors |  |  | GEO accession |
| --- | --- | --- | --- | --- | --- | --- | --- | --- |
|  |  |  |  |  | STAT | IRF | ETS |  |
| Type I | IFNa | 6h and 12h | 500U/ml | HT1080 | STAT1 (1), STAT2 (6), STAT5B (22) |  | ELF1 (14) | GSE31019 |
|  |  | 6h | 500U/ml | SKOV3 | STAT1 (2), STAT2 (11) | IRF7 (5), IRF1 (13), IRF2 (18) | ETV6 (20), ELF1 (30) | GSE31019 |
|  |  | 6h | 10U/mL | Primary Hepatocytes | STAT2 (12), STAT1 (24) | IRF7 (4), IRF9 (7), IRF1 (17), IRF6 (29) | ETV7 (1), ETV6 (27), ELK4 (28) | GSE31193 |
|  |  | 24h | 10U/mL | Primary Hepatocytes | STAT2 (1), STAT4 (7), STAT1 (9) | IRF7 (3), IRF9 (11) | ETV7 (4), ELK4 (17) | GSE31193 |
|  |  | 10 h | 1000 U/ml | Fibroblast | STAT1 (8), STAT2 (24) | IRF9 (7) | ELF1 (32) | GSE67737 |
|  | IFNa2 | 6 h | 1000 U/ml | Keratinocyte | STAT1 (6), STAT2 (9), STAT5A (32) | IRF7 (20), IRF1 (21), IRF6 (25), IRF2 (28) | ETV7 (11), ELF1 (26) | GSE124939 |
|  |  | 18 h | 1000 U/ml | Primary macrophage | STAT3 (9), STAT1 (13) | IRF1 (26), IRF7 (27) |  | GSE30536 |
|  | IFNa2b | 2 h | 1000 U/ml | EBV-transformed B lymphoblastoid cell lines |  |  | ELF1 (2) | GSE117637 |
|  |  | 2 h | 1000 U/ml | Fibroblast |  | IRF2 (6) | FLI1 (29) | GSE117637 |
|  | IFNa6 | 6 h | 1000 U/ml | Keratinocyte | STAT1 (14), STAT2 (17) | IRF7 (11), IRF1 (28), IRF6 (30) | ETV7 (9), ELF1 (32) | GSE124939 |
|  | IFNb | 6 h | 1000 U/ml | Keratinocyte | STAT1 (8), STAT2 (15) | IRF1 (5), IRF7 (6), IRF2 (19), IRF6 (20) | ETV7 (1), ELF1 (18) | GSE124939 |
|  |  | 10 h | 1000 U/ml | Fibroblast | STAT1 (10), STAT2 (30) | IRF9 (8) | ETV7 (34), ELF1 (35) | GSE67737 |
| Type II | IFNg | 6 h | 5 ng/ml | Keratinocyte | STAT3 (13), STAT1 (23), STAT2 (24) | IRF2 (1), IRF1 (5) | ETV7 (8) | GSE124939 |
|  |  | 20 h | 10 ng/ml | Monocyte-derived macrophages | STAT1 (2), STAT2 (25) | IRF1 (1), IRF9 (31) | ETV7 (3) | GSE79077 |
|  |  | 6 h | 20 ng/mL | Monocytes | STAT2 (9), STAT1 (11) | IRF1 (3), IRF9 (15), IRF7 (22) | ETV7 (1) | GSE36537 |
|  |  | 18 h | 20 ng/mL | Monocytes |  |  |  | GSE36537 |
|  |  | 18 h | 20 ng/mL | Monocyte-derived macrophages | STAT1 (17), STAT2 (34) | IRF1 (2), IRF7 (35) | ETV7 (1) | GSE36537 |
|  |  | 24 h | 20 ng/mL | keratinocytes | STAT3 (3) | IRF1 (1), IRF2 (10), IRF7 (14) | ETV7 (2) | GSE12109 |
|  |  | 24 h | 100 U/ml | Peripheral blood derived macrophages | STAT3 (4) | IRF7 (3), IRF1 (8), IRF9 (26) | ETV7 (2), ELF4 (27) | GSE11886 |
|  |  | 10 h | 1000 U/ml | dermal fibroblast | STAT1 (8), STAT3 (23) | IRF1 (4), IRF9 (10) | ETV7 (11) | GSE67737 |
|  |  | 3 h | n.a. | Monocyte-derived macrophages | STAT2 (12), STAT6 (30) |  | ETS2 (5), ELK1 (24) | GSE130567 |

\* Three families of transcription factors were assessed for 6 datasets, where cells were stimulated by different interferons. Only TFs in STAT, IRF, and ETS family ranked in top 35 were shown in “Families of transcription factors” columns as a format of “Regulator (ranking)”. The best combination of
parameters (i.e., COEN network and GSEA enrichment method) identified in Table 1 were used in the analysis.

A recent work has shown that ELF1 (a member of ETS transcription factor family) is induced by IFN, but does not feed-forward to induce interferons, and transcriptionally programs cells with potent antiviral activity ^43^. Interestingly, ELF1 was identified by *RegEnrich* as one of the key regulators in most of the type I IFN stimulation experiments (**Table 2**). We further investigated whether any other members of the ETS transcription factor family were also identified by *RegEnrich*. Interestingly, we found an ETS transcription factor family member, ETV7, in the lists of top regulators from more than half of type I IFN stimulation experiments and from almost all type II IFN stimulation experiments. A more recent study shown that ETV7 preferentially targeted a subset of antiviral ISGs that were crucial for IFN-mediated control of viruses, such as influenza and SARS-CoV-2 ^44^.

Different cells may respond differently to IFN stimulation with different durations. We further investigated the common regulators involved in IFN stimulation among different cells. Thus, we summarized the most common regulators within type I and type II IFN stimulation experiments. It was shown that the ISGs of type I IFNs were strongly regulated by STAT family, TRIM family, IRF family, ETS family, SP100/SP140 family (transcriptional coactivator of ETS family TFs), etc. Similarly type II IFN ISGs were largely regulated by STAT family, IRF family, ETS family, MCM family, SP100/SP140 family, etc. (**Table 2** and **Fig. 5**) The most commonly identified regulator of type II IFN ISGs was MHC class II transactivator (CIITA), which has been very recently shown with potential to induce cell resistance to Ebola virus and SARS-CoV-2 ^45^. Altogether, these results suggest that *RegEnrich* successfully identified key regulators related to IFN signaling in IFN stimulation experiments.

**Fig. 5.**
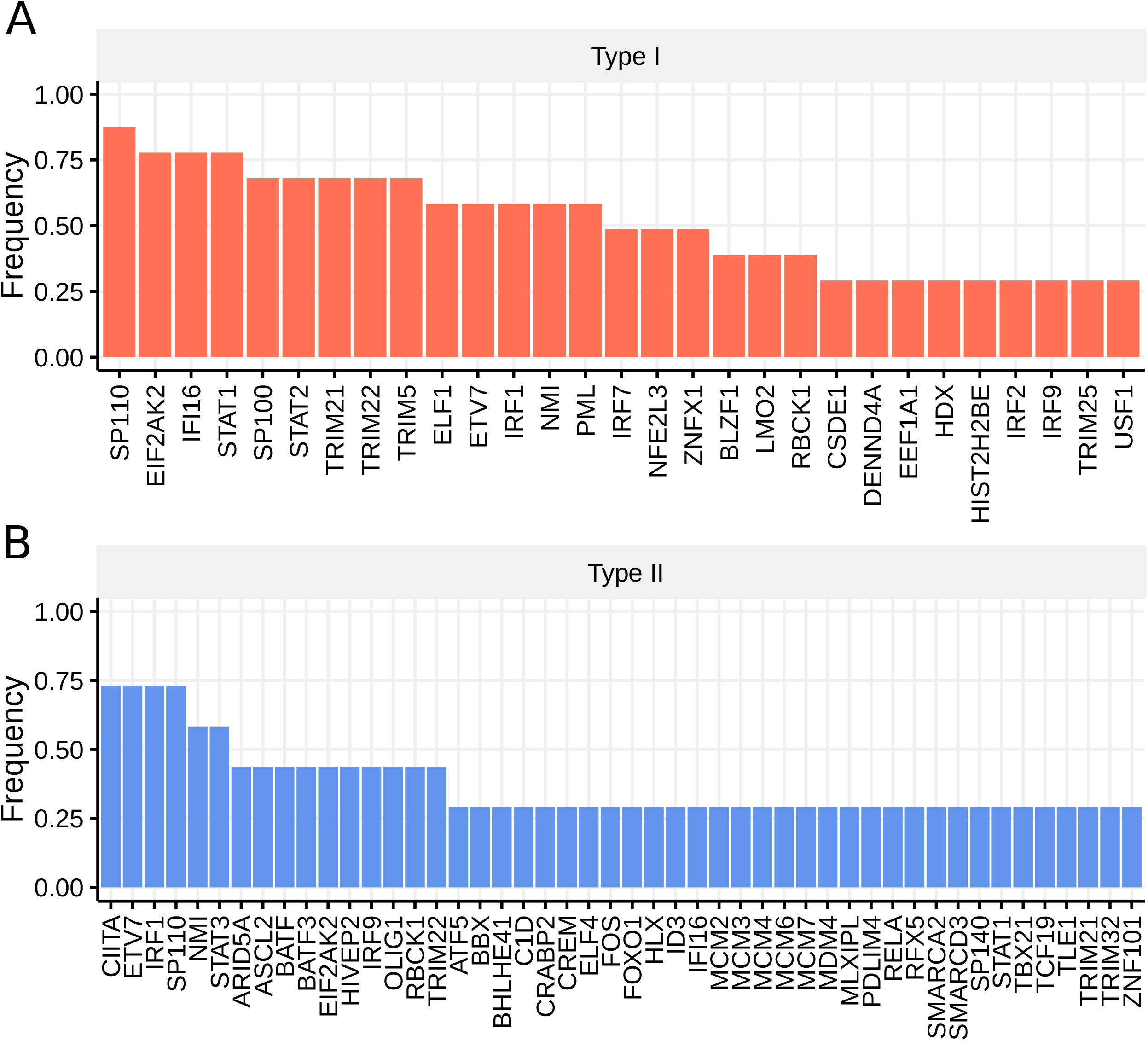
The genes consistently identified as key regulators in **(A)** type I interferon stimulation and **(B)** type II interferon stimulation datasets that were shown in **Table 2**. The top 35 regulators in each dataset were included as key regulators, and only the regulators identified in more than 25% of datasets were shown.

## 4. Discussion

High throughput technologies like microarray, RNA-seq and protein mass spectrometry offer easy, fast, and affordable profiling of the gene/protein expression. These technologies generate massive data facilitating us to study the alterations in gene/protein expression, thereby helping us to identify the biomarkers for diseases and for other biological states. However, it is still challenging to predict which genes play the major roles in these biological contexts. To address this problem, we developed *RegEnrich*, an open-source R/Bioconductor package integrating differential gene expression analysis, network inference, enrichment analysis and regulator ranking. *RegEnrich* is able to identify the key regulators by providing gene/protein expression data from multiple high throughput technologies. We benchmarked the speed and maximum usage of network inference methods in the *RegEnrich*, which shows that the COEN method runs much faster than random forest methods when number of genes is below 20,000, and the speed of multi-threaded random forest version is significantly improved. Traditionally, COEN is considered as a method for depicting linear relation between genes while the random forest for a non-linear relation. Strikingly, in the Lyme disease transcriptomics dataset the results from these two methods were consistent. This might be because COEN methods re-evaluate the edges weights by considering the information of neighbor nodes, as a result, such a network was constructed not only based on a linear relationship.

Since hub nodes have been found to be important in many networks, hub genes that are defined by gene regulatory network properties, are also expected to be crucial in biology as well and have drawn much attention over last decade ^46^. We compared the hubs identified by the network properties, degree and closeness, with the key regulators by *RegEnrich*. Interestingly, only a very small number of the key regulators from either *RegEnrich* or VIPER were hubs (**Fig. 4A, B**). One possible reason is that these hub genes are so important in maintaining the major functions of cells that too strong perturbations of these hubs could be fatal for cells ^47^. Therefore, these hub genes are not necessarily the key gene expression regulators in a specific context. For example, Gaiteri *et al*. showed that differentially expressed genes primarily reside on the periphery of co-expression networks for neuropsychiatric disorders such as depression, schizophrenia and bipolar disorder ^48^.

As one easy-to-use software/package whose functionality was similar to *RegEnrich*’s, VIPER R package was used to compare with *RegEnrich* to find the final key regulators. According to the instruction of VIPER package on Bioconductor, VIPER needs an ARACNE network to perform the analysis. Such network is generated by an independent package, such as *minet* ^49^, *GPU-ARACNE* ^50^, and *ARACNe-AP* ^16^. These packages either are not R packages or fail to construct the network when the sample size is small. In contrast, *RegEnrich* is an all-in-one package and a detailed tutorial document is provided along with *RegEnrich*, which facilitates users to use it more easily. More importantly, *RegEnrich* can identify the key regulators whose expression and their targets’ expression correlate with the experimental phenotypes. In addition, different from the VIPER package, *RegEnrich* is able to find key regulators not only between two conditions but also in a time series experimental setting. Since *RegEnrich* is modular and is intended to be a flexible pipeline, we allow users with options to provide custom regulator lists, and options for multiple methods at different steps in case needed.

Every method has its own disadvantages, *RegEnrich* is no exception. Because the final key regulators are dependent on the gene regulatory network, the major disadvantage of *RegEnrich* is that, presently, the date-driven network is inferenced by only gene/protein expression data, which is also a disadvantage of ARACNE + VIPER pipeline. Glass et al. has shown that integration of protein-protein interaction, protein-gene interaction and gene expression can increase the accuracy of regulatory network inference ^51^. Currently, we provide an option for the users to provide their own gene regulatory network, which can be derived from other epigenetic datasets such as ChIP-seq, ATAC-seq data, protein-protein interactions etc., thus, granting *RegEnrich* an ability to integrate multi-omic data.

By analyzing datasets of dozens of IFN-stimulation experiments, *RegEnrich* identified STAT and IRF transcription factor family members, including STAT1, STAT2, IRF9 and IRF7, which have been extensively shown to play important roles in IFN signaling pathways ^41,42^. Meanwhile, *RegEnrich* also identified several ETS transcription factor family members, such as ELF1 and ETV7, as key regulators in IFN signaling. Interestingly, ELF1 transcriptionally program cells with potent antiviral activity, and ETV7 targeted antiviral ISGs crucial for IFN-mediated control of virus including influenza and SARS-CoV-2 ^43,44^. These antiviral activities are typical the fundamental roles of IFN in innate immunity. By analyzing the most commonly top-ranked regulators, *RegEnrich* gave a list of candidate key regulators, such as CIITA and SP100/SP140 family members. Given CIITA has been recently reported with antivirus ability ^45^, further study may be carried out to investigate the anti-virus potential of SP100/SP140 family members, such as SP100 and SP110, which might facilitate the mechanistic studies of IFN-ISG signaling and ultimately drug development.

Recently, using RegEnrich pipeline, we predicted a network of key regulators that leads monocyte-derived dendritic cells (moDCs) to differentiate into a different trajectory upon CXCL4 stimulation, compared to the moDCs without CXCL4 stimulation. We also experimentally validated *RegEnrich* pipeline’s prediction by silencing one of the top-ranked regulators in the predicted network, i.e., CIITA ^52^. More recently, we studied the mechanism of human T regulatory (Treg) cells programming under inflammatory conditions. Using RegEnrich, we predicted a network of key regulators important for effector Treg differentiation, including the vitamin D receptor (VDR), which is further validated by H3K27ac and H3K4me1 ChIP-seq experiments ^53^. These two independent experimental studies support that RegEnrich is able to accurately rank the key gene regulators that are mechanistically involved in immune cell development and functions.

## 5. Conclusions

Understanding the key regulators between different biological states is essential for gaining mechanistic insights, designing functional experiments, and rational drug development. To this end, in this paper we presented *RegEnrich*, a Bioconductor R package for inference of key regulators in biological conditions. There are four major steps to obtain the list of key regulators in *RegEnrich*, i.e., differential expression analysis, regulator-target network inference, enrichment analysis and regulator ranking. For differential expression analysis, the methods in DESeq2 and limma packages are provided, which grants *RegEnrich* ability to predict the key regulators not only for gene expression data of two conditions but also for time series data. Meanwhile, two regulator-target network inference methods, WGCNA and random forest, are provided, which allows the network not only contains the linear information but also include non-linear relationship between genes. FET and GSEA algorithms are optional for user to perform enrichment analysis. *RegEnrich* can identify the key regulators whose expression and their targets’ expression correlate with the experimental phenotypes. Using datasets from gene silencing studies, *RegEnrich* using GSEA method performed the best to retrieve the key regulators and outperformed VIPER package. Further, by analyzing dozens of *in vitro* interferon-stimulation gene expression datasets, *RegEnrich* identified not only IRF and STAT transcription factor families played an important role in cells responding to IFN, but also several ETS transcription factor family members, such as ELF1 and ETV7, are highly associated with IFN stimulations. Above all, *RegEnrich* can accurately identify, in a data-driven manner, key gene regulators from the cells under different biological states, which can be valuable in mechanistic studies of cell differentiation, cell response on drug stimulation and disease development, ultimately in drug development.

## Acknowledgements

We appreciate the critical discussions within CoSI group, UMC Utrecht, which significantly improved this manuscript.

## Authors’ contributions

WT and AP conceptualized the idea and designed the methodology. WT designed the algorithmic solutions and wrote the code. AP supervised the research. AP and TRDJR provided the resources for the research. WT, AP and TRDJR wrote the original draft, and reviewed and edited the manuscript. All authors contributed to conceiving the idea. All authors read and approved the final manuscript.

## Ethics approval and consent to participate

Not applicable

