## Supplementary for "RegEnrich: An R package for gene regulator enrichment analysis reveals key role of ETS transcription factor family in interferon signaling"

### Supplementary Materials

Table S1. Memory usage and time consumption by RegEnrich analyzing different size of data.

| Time consumption in second (4 cores, GRN) | | | Time consumption in second (1 core, GRN) | | | Time consumption in second (4 cores, COEN) | | | | Time consumption in second (1 core, COEN) | | | | |
|---|---|---|---|---|---|---|---|---|---|---|---|---|---|---|
| Number of samples | | | Number of samples | | | Number of samples | | | | Number of samples | | | | Number of genes |
| 200 | 100 | 50 | 200 | 100 | 50 | 100 | 50 | 20 | 10 | 100 | 50 | 20 | 10 | |
| 78463.2 | 29658.8 | 10822.4 | 211594 | 77376.1 | 31182.6 | 87771.6 | 92128.7 | 100301.6 | 92397.6 | 91148.4 | 87938.5 | 89958.2 | 92357.8 | 40000 |
| 77735.5 | 28736.7 | 10070.1 | 170517.9 | 62417.9 | 25241.3 | 55702.7 | 55819.5 | 55736 | 55876.9 | 55724.6 | 55878.4 | 55771.8 | 55950.3 | 35000 |
| 64384.9 | 23591 | 9645.4 | 154590.9 | 53700.8 | 22305.2 | 36901 | 36944.9 | 36863.3 | 36883.6 | 37083 | 36862.8 | 36861.6 | 36864.6 | 30000 |
| 51112 | 18641.1 | 7463.6 | 130287.2 | 47389.4 | 18640 | 22061.3 | 22051.4 | 22073.2 | 23654.5 | 23688 | 23678.9 | 23672.9 | 23663.5 | 25000 |
| 42765.5 | 14317.7 | 5835.8 | 112109.5 | 36372.9 | 14645 | 11876.2 | 11893.1 | 12170.4 | 12150.9 | 12273.5 | 12182.5 | 12154.6 | 12165.6 | 20000 |
| 32415.3 | 12081.5 | 4525 | 82561.1 | 27112.2 | 10925 | 5205.8 | 5187.8 | 5182.5 | 4121.3 | 4172.3 | 5188.2 | 5170.5 | 4198.7 | 15000 |
| 25017.1 | 9251.3 | 3382.3 | 58231.3 | 22734.8 | 8330.8 | 2272.1 | 2706.8 | 2704.5 | 2705.2 | 2712 | 2700.9 | 2703.7 | 2703 | 12000 |
| 20161.2 | 8085.8 | 2951.1 | 52217.1 | 20132.2 | 7864.3 | 1634.3 | 1398.2 | 1400 | 1401.5 | 1356 | 1623.2 | 1628.3 | 1628.4 | 10000 |
| 15879.9 | 6578.8 | 2439.4 | 37921.2 | 15003 | 5815.2 | 770.6 | 802.2 | 803 | 685.8 | 795.3 | 880.8 | 846.5 | 847.3 | 8000 |
| 13924.8 | 5156.2 | 2050.1 | 35534.2 | 13590 | 5153.7 | 610.3 | 607.8 | 609 | 610.9 | 608.8 | 605.7 | 608.4 | 595.3 | 7000 |
| 12290.9 | 4353.3 | 1402.1 | 30828.4 | 11970.6 | 3496.7 | 406.8 | 407.4 | 407.8 | 408.9 | 408.5 | 405.7 | 406.6 | 407.7 | 6000 |
| 10828.4 | 3854.1 | 1458.7 | 27986.3 | 9436.9 | 3925.5 | 257.2 | 256.5 | 255.7 | 256.4 | 255.7 | 252 | 250.7 | 260.8 | 5000 |
| 8277.3 | 3061.8 | 1183.8 | 15518.4 | 8074.4 | 3040.7 | 145.5 | 149.1 | 149.6 | 156.6 | 156.1 | 132.6 | 165.7 | 146.5 | 4000 |
| 6223 | 2257.3 | 790.6 | 15395.7 | 5940.3 | 2250.9 | 84.7 | 86.7 | 82.7 | 80.6 | 71.7 | 81.7 | 85.5 | 82.4 | 3000 |
| 3855.5 | 1414.3 | 533 | 10747 | 3905.6 | 1387.2 | 35.6 | 36.7 | 36.6 | 41.6 | 39.3 | 39.1 | 37.3 | 38 | 2000 |

Table S2. Top 35 regulators using FET and GSEA methods of RegEnrich in Table 1.

| GEO accession | Silencing technology | No. of samples | Cell line | Silenced gene(s) | Ranking | | Genes | |
|---|---|---|---|---|---|---|---|---|
| | | | | | RegEnrich (FET) | RegEnrich (GSEA) | RegEnrich (FET) | RegEnrich (GSEA) |
| GSE19114 | shRNA | 44 | BTIC | STAT3 | 2 | 1 | MED6 STAT3 TCEA2 HIF1AN TCF3 LCOR ELF4 E2F6 STAT2 SNAPC3 KLF13 CDC5L ELF2 TAF1B POLR2G SND1 TAX1BP3 MTF2 CRY1 LZTS1 ETV6 TOX4 PCGF2 GTF2E2 RERE ZNF35 GTF2A2 DR1 NCOA3 BCL6 SMARCA5 MIF THAP7 E2F4 ELK1 | STAT3 MED6 STAT2 LCOR CDC5L CRY1 ELF2 HIF1AN E2F6 TCEA2 ELF4 POLR2G KLF13 BCL6 LZTS1 DR1 SMARCA5 TCF3 SNAPC3 SND1 TAF1B GTF2A2 BTG1 TOB1 TAX1BP3 GTF2B TOX4 ETV6 ID3 DRAP1 ERF SIRT7 TRIP13 MTF2 PRRX1 |
| GSE17172 | shRNA | 9 | ST486 | FOXM1 | 2 | 1 | FOXN3 FOXM1 TBPL1 FOXG1 ESRRA ETS1 AATF EVX1 AFF1 RUNX3 NDN ARID3A APBB1 PIAS4 E2F5 CREB3L2 APEX1 SMAD4 XBP1 SOS1 ASCC3 HIST1H2BJ HOXC6 MTF2 PROP1 DR1 EHMT2 TAF1C GTF2A1 NFX1 TCF3 POU6F1 PLAGL1 E2F4 NKX2-1 | FOXM1 CREB3L2 EVX1 PIAS4 FOXN3 XBP1 HIST1H2BJ TAF1C E2F5 FOXO1 NDN ARID3A SOS1 MMS19 TBPL1 FOXG1 NR1D2 AFF1 ETS1 DAB2 HIST1H2BD CREB3 NFYA EHMT2 ASCC3 KLF12 SMARCC1 APBB1 SMAD4 TARBP2 FOXJ3 AATF PROP1 PWP1 DLX4 |
| GSE19114 | shRNA | 12 | SNB19 | STAT3 | 14 | 5 | ELF4 VEZF1 HMGB2 TEAD2 MED6 SP2 NFE2L1 TFDP3 TP53BP1 CCDC6 SOX18 ZIC2 STAT2 STAT3 VHL SNAPC3 RXRA UPF1 LZTR1 KLF13 ZEB2 KLF2 ID3 E2F6 ARID5B CEBPA LCOR TCEA2 HOXB5 ELF5 TSC22D1 ETV6 PCBD2 FOXO1 NR2C2 | VEZF1 SNAPC3 SOX18 TEAD2 STAT3 ID3 LZTR1 KLF2 VHL PRRX1 ETV6 STAT2 ZIC2 NFE2L1 E2F6 BIN1 ZNF165 MED6 ELF4 ARID5B CCDC6 UHRF1 PHF10 SAFB2 ZNF467 TAX1BP3 BMI1 TSC22D1 JAZF1 ZNFX1 RXRA BBX NCOR2 KLF9 SIX5 |
| GSE2350 | siRNA | 8 | Burkitt lymphoma | BCL6 | 32 | 11 | PDLIM1 SSBP2 E2F5 POU2F1 MYBL1 HESX1 NR3C1 IFI16 MXD1 POLR2J NR5A2 CUX2 ZNF267 MNT ZNF282 TCEB1 NEUROG3 TCEB3 PNN CRY1 TSC22D1 COIL EN2 IKBKB ELF1 RUNX2 SREBF1 HLTF TCF12 ZBTB22 ENO1 BCL6 JARID2 TCF20 TCEA1 | PDLIM1 HESX1 SSBP2 CUX2 ELF1 NR5A2 CRY1 COIL TSC22D1 MYBL1 BCL6 ZNF510 IKBKB E2F5 NEUROG3 MXD1 POU2F1 MNT ZNF282 ZNF267 CEBPG TCEB3 RUNX2 MAFF ZNF175 AHR LHX2 RFXAP ZNF354A NAB1 RNF4 EN2 POU6F2 FOXK2 ID3 |
| GSE19114 | shRNA | 12 | SNB19 | CEBPB | 50 | 24 | JAZF1 PHF10 HIC2 SMAD5 POLR2C HMGN1 TOB1 ELL2 HIST1H2BD ID1 MED30 VEZF1 CNOT7 SCMH1 EIF2AK2 PRDM1 CRABP2 PCGF6 MED4 ZNF175 ATF5 CITED2 HIPK2 ZNF263 ID3 SAFB2 FBXO7 TCF12 E2F6 GMEB2 FHL2 ARID3A SNAI2 CREG1 NCOA1 | PAPOLA SOX18 SUPT16H CREB3L2 PARP1 CDCA7L GTF2F2 NCOA3 PRRX1 SNAPC3 FOXO3 PCGF6 MED20 CEBPG PER2 SMAD5 ID3 ZBED1 JAZF1 SMAD6 ZFHX3 HMGN1 SOX21 CEBPB CITED2 EPAS1 ERCC3 HIST1H1C PHF10 HIC2 VEZF1 MCM3 RFX5 MAFG TCF12 |
| GSE19114 | shRNA | 44 | BTIC | STAT3 & CEBPB | 2 & 285 | 1 & 957 | TCEA2 STAT3 SNIP1 TOB1 NCOA3 CITED2 PRDM4 RFXAP PHF10 MAFG E2F5 ZNF75A MED8 MED30 SUPT16H KNTC1 TOE1 RFX5 SERTAD2 RNF14 SUZ12 MTF2 PAPOLA HES1 FOS TAX1BP3 NDN EP400 HMG20A POLR2C TRIM5 NKX3-1 ID1 RB1 CENPK | STAT3 NCOA3 MED8 E2F5 SNIP1 SUPT16H KNTC1 CITED2 PHF10 RNF14 TOE1 PRDM4 MTF2 EP400 RFXAP PAPOLA TOB1 HES1 FOS NFYC MED30 POLR2C ZNF75A HMG20A RFX5 SERTAD2 MED6 POLR2J TCEA2 ZNF451 MED15 ZBTB43 VEZF1 IRF9 MLLT10 |
| GSE19114 | shRNA | 12 | SNB19 | STAT3 & CEBPB | 365 & 6 | 38 & 11 | VEZF1 ID3 ID1 PHF10 JAZF1 CEBPB PRRX1 TOB1 ZNF467 ELL2 MED30 ZNF263 MXD3 RNF14 HCFC1 PRDM1 NFKBIA PHF21A FHL2 ATF5 FOSL1 CRABP2 SAFB2 NKX2-2 SOX13 HMG20B NCOA1 ZNF35 HDAC4 ASCC2 POU3F2 HDAC1 ZNF451 SOX18 ATF4 | SOX18 PRRX1 VEZF1 CREB3L2 SUPT16H PAPOLA ID3 PARP1 ZFHX3 NCOA3 CEBPB ID1 FOXO3 CDCA7L ZBED1 MED20 SMAD5 PHF10 EPAS1 PCGF6 HIC2 SOX21 JAZF1 CEBPG PER2 MED30 SOX9 ID2 TCF4 SMAD6 GTF2F2 HMGN1 TOB1 ELF1 MCM3 |
| GSE51978 | shRNA | 9 | IMR32 | CHAF1A | 10 & 29* | 15 & 46* | (ZFP36L1 HR KAT2B ZBTB24 RREB1 LMX1A ABT1 PLAGL1 ETV3 CHAF1A ARID1B PRDM16 SOX15 ATF6B RXRG EGR1 SALL1 MYB VHL MAPK14 STAT5A MBD2 SNAPC1 RAI14 CEBPD HIST1H2BD POLR2L ZNF274 LHX8 NRIP1 TRPS1 SS18L1 PRPF4B MDM4 POLR1B) & (RREB1 PLAGL1 RAI14 ZFP36L1 DDB2 RARB LMX1A MLX HR CEBPD MAPK14 ARID1B NRIP1 LHX8 PRPF4B MBD2 NR4A3 APC KAT2B SNAPC1 HIF1A ATF6B SPI1 TBX1 ZNF146 NHLH1 SSRP1 INSM1 CHAF1A SALL1 PRDM16 NEUROD1 FOXC1 MAFG SUZ12 ) | (ZFP36L1 HR PRDM16 KAT2B SOX7 PLAGL1 ZBTB24 RREB1 LMX1A ABT1 TRPS1 ETV3 FOXO6 DTX1 CHAF1A ARID1B EGR1 FOXD2 BHLHE23 VSX1 SOX15 FOS FOSL1 SIX4 ATF6B TAF4 RXRG RAI14 ELL2 HIST1H2BD ATF2 POU1F1 SALL1 DLX3 ASB9) \| (ZFP36L1 HR RREB1 PLAGL1 PRDM16 NHLH1 RAI14 TBX1 GMEB1 DDB2 NR4A3 RARB LMX1A FOXO6 HIST1H2BJ MAFG MLX SOX7 ZSCAN4 ARID1B SNAPC1 POU2F3 VGLL1 KAT2B ZNF287 SPI1 ELF2 NEUROD6 ELL2 ZNF488 SALL1 CEBPD FOXC1 TRPS1 HOXD4) |
| GSE19114 | shRNA | 44 | BTIC | CEBPB | 913 | 793 | NCOA3 SNIP1 PAPOLA MED30 ELF4 ELF2 TRIM5 MAFG NFYC E2F5 MED8 ERCC3 RFXAP SIX4 POLR2C NDN CITED2 SNAPC3 PHF10 SERTAD2 MED20 SUZ12 ATF2 GLI3 TRIB3 SCMH1 E2F6 EP400 DAB2 CENPK GTF2B PRDM4 CREBBP POLR2J TOB1 | NCOA3 E2F5 PAPOLA ELF2 NFYC MED8 SNIP1 MED30 MED20 E2F6 SERTAD2 POLR2C GTF2B EP400 SCMH1 SIX4 PRDM4 RFXAP GLI3 DAB2 ERCC3 HBP1 TOB1 ELF4 CENPK MAFG TRIM5 ZNF408 RFX5 ZNF384 NDN RB1 PARP1 ZNF559 TRIM21 |

#### Supplementary figures

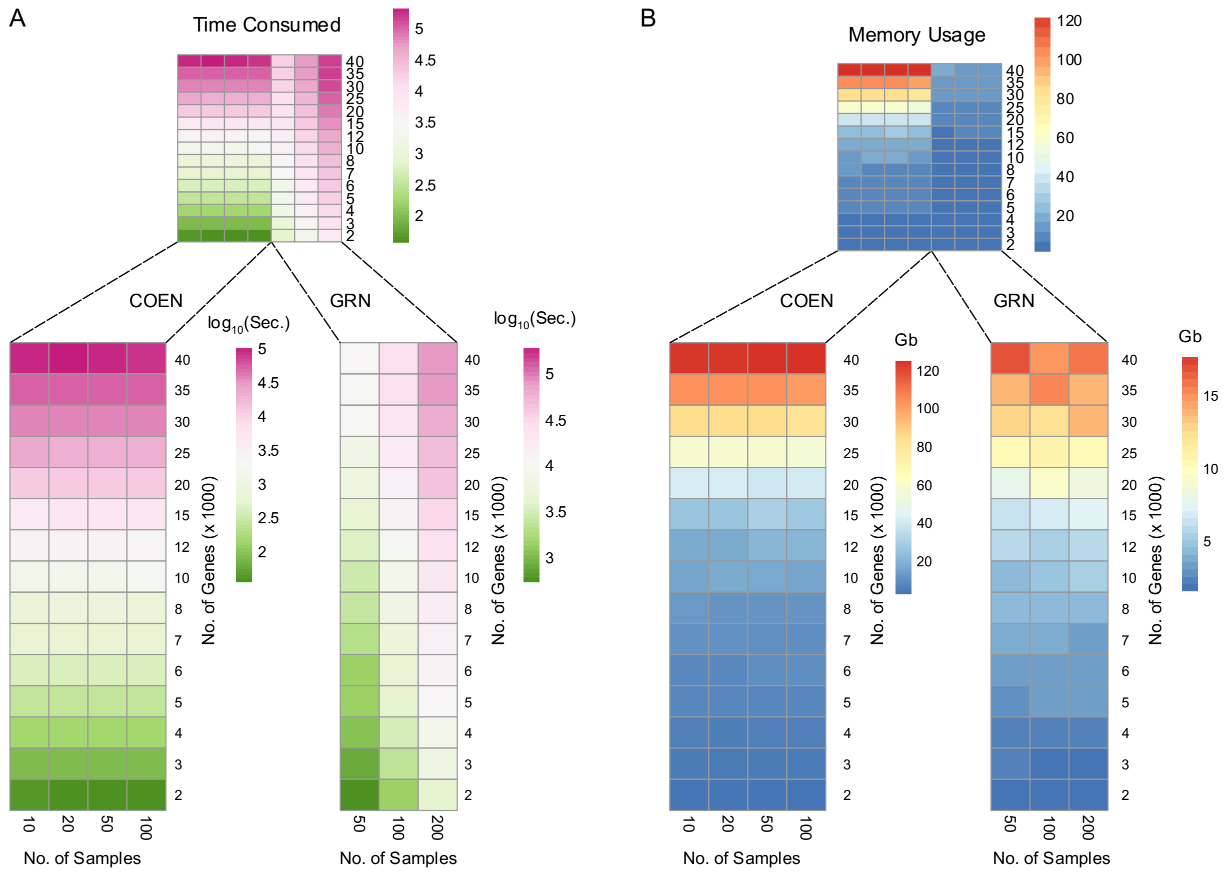


**Fig. S1**. Time consumption and memory usage by *RegEnrich*. With four CPU cores, (A) the time consumed, and (B) maximum memory used, by *RegEnrich* when analyzing a gene expression dataset with different number of genes (ranging from 2,000 to 40,000) and different number of samples (ranging from 10 to 100 and from 50 to 200 for WGCNA and random forest network, respectively).

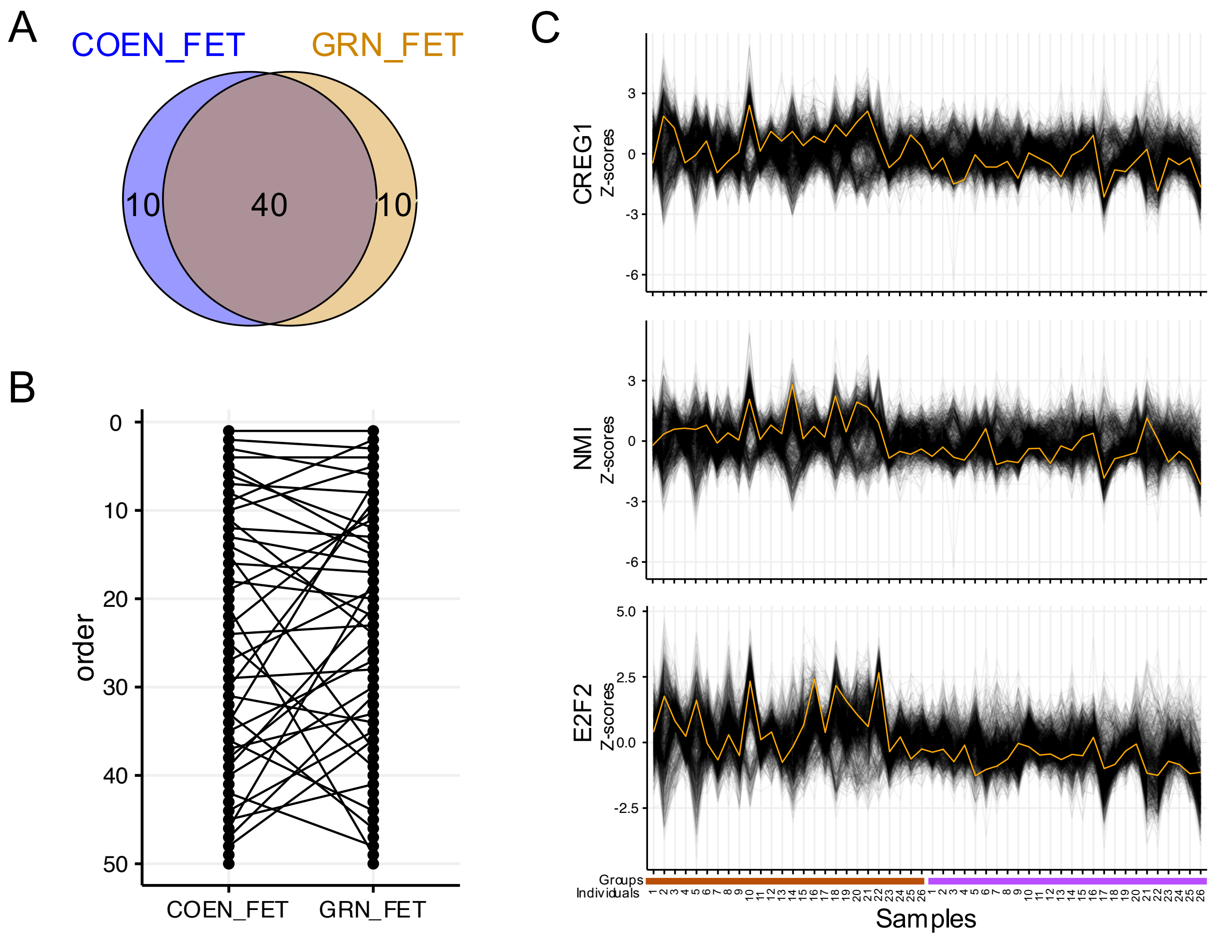


**Fig. S2**. Comparison of top 50 key regulators from different network inference methods in *RegEnrich* package. (**A**) Venn diagram of key regulators from WGCNA network and random forest–based network. (**B**) Plot of regulator orders: the regulators identified by each package are ordered by their corresponding importance scores, and the same regulators are linked by lines. (**C**) The expression pattern of top three key regulators and their targets identified from random forest–based network. Yellow lines are the normalized expression of regulators and grey lines are of corresponding targets.
